# Central Nervous System Control of breathing in Natural Conversational Turn-Taking

**DOI:** 10.1101/2024.07.17.603521

**Authors:** Camilla Di Pasquasio, Lila De Pellegrin, Arthur Pineaud, Antonin Marty, Thierry Chaminade

## Abstract

Discussion is a fundamental social activity requiring coordination of speech between interlocutors. Speech production is a complex human behaviour that involves several anatomo-physiological processes, including inspiration and expiration. The aim of the present study is to investigate the neurophysiological underpinnings of speech-related respiration events in conversational turn-taking. We made use of an existing corpus of natural conversations between a participant and its interlocutor (Human or Robot) focusing on synchronised (1) behavioural (conversation turn-taking), (2) respiratory (maxima of inspiration) and (3) neurophysiological (fMRI) data. Precisely timed conversation transcripts from 25 participants were used to categorise breathing maxima based on their timing relative to the participant’s speech onset. In agreement with the literature, the closest respiration time maximum to each speech turn occurred on average 200 ms prior to speech onset. The fMRI second-level contrast (*p_FWE_* < 0.05, extend k > 5 cm^3^) Resp+ (maximum respiration associated with speech) *versus* Resp-, exclusively masked to exclude speech related areas, revealed bilateral activations in the central sulcus, the brainstem and the cerebellum. The brainstem cluster comprises respiratory pattern generators, possibly the preBötzinger complex, that need to be inhibited to enslave breathing to speech production and not physiological needs, while the central sulcus cluster is likely to be located in the postcentral primary sensory cortex receiving upper torso inputs indicating that lungs are filled, and the cerebellum clusters could play a role in the timing of speech onset, 200 ms after a respiration maximum. These results show how cortical, cerebellar and brainstem coordinated control of breathing during conversational turn-taking is part of the intricate physiological mechanisms that contribute to natural communication dynamics.

## Introduction

Language is a unique hallmark of human cognition, and conversations, from everyday chit chat to specific discussions, play a widely acknowledged function in social interactions (Hudson, 1996). Considering the need to investigate neural correlates of “*real-time social encounters in a truly interactive manner*” (Schilbach et al., 2013), we present here an investigation of the physiology associated with the control of respiration in natural conversations.

### Conversation, a social implementation of speech

Discussion is characterised by the exchange of spoken remarks between two or more interlocutors. This trivial everyday activity plays a very important social role in our societies. To enable this interaction, one speaker must be able to perform a phonatory exhalation to express its remarks, as well as to control the turn of speech between himself and his interlocutor. Smooth conversation entails accurate prediction of the turn-taking time, i.e. the time when one interlocutor stops speaking and therefore leaves the floor open for the other speaker. Fluid transitions between speakers typically occur within a temporal window of a few hundred milliseconds (Torreira et al., 2015). Reducing the turn-taking time increases the amount of factual information that can be exchanged between speakers in a limited amount of time. Note that a study of a variety of languages shows a universal tendency to avoid overlapping conversations and minimise silences between speech turns (Stivers et al., 2009).

Speech is a complex behaviour that requires precise coordination of multiple anatomo-physiological processes. During phonatory exhalation, a series of events are triggered to enable sound production. The vocal folds and soft palate open during this phase, allowing air from the lungs to enter the oral cavity. The subglottic pressure generated by this air flow concentrates under the vocal cords. The latter, in an open abducted position, rub against each other and vibrate, producing the phonemes that form the basis of spoken language. This phenomenon is closely linked to laryngeal pressure, a force resulting from the interaction between the vocal folds and subglottic pressure. This laryngeal pressure is essential to distinguish breathing from vocalisation, underlining the importance of muscular control in the process. The diversity of phonemes emitted then depends on the positioning of the oral and nasal cavities, contributing to the richness of speech (Demartsev et al., 2022). This control is exercised by the airway muscles, often referred to as a “valve”, as well as by the lower tract, acting as a thoracic bellows. These muscular components, represented in the dorsal primary motor cortex, contribute to the precise regulation of breathing required for sound production (McKay et al., 2003). In contrast, articulation is associated by the involvement of the ventral primary motor cortex (Pulvermüller et al., 2006), that have direct access to the motor neurons that control the muscles involved in speech, such as those of the face, mouth, tongue, larynx and pharynx (Holstege & Subramanian, 2016). Altogether, these results confirm a dissociation between the functions of dorsal and ventral motor cortices in human speech (see also Price, 2012), that would correspond to the control of respiration and articulation, respectively.

### Central nervous system control of speech

The neural bases of articulation in speech have been - and still are - widely investigated. Patient Louis Victor Leborgne, studied by neurosurgeon Paul Broca, was unable to express other syllables than Tan and presented a lesion in the left inferior prefrontal cortex. The link between *a* (absence) - *phasia* (speech) and a localised brain lesion is at the origin of neuropsychology, the investigation of causal links between lesions in specific brain regions and associated behavioural deficits. In contrast, little is known about the neurophysiological control of breathing in speech, despite its importance in particular in conversations. Preparing to speak indeed involves subtle coordination of interlocutors’ breathing cycles. Speakers plan their speeches by synchronising their inspirations with the end of questions, thus establishing a dynamic temporal relationship between turn-taking and breathing (Rochet-Capellan & Fuchs, 2014). Other investigations revealed that inspirations before long responses cluster around the end of the question, while inspirations before short responses show greater dispersion and at earlier timings, suggesting a distinction between inspirations specifically intended for speech and those more related to physiological needs (Torreira et al., 2015). Breathing indeed adapts to communication needs during smooth conversations, underlining the dynamic and adaptive aspect of the process.

While the central nervous system control of respiration in natural conversations remains unknown, neuropsychological literature reports a significant number of cases of dissociation between automatic and voluntary breathing. A patient study on a case of severe respiratory apraxia resulting from supranuclear palsy revealed an inhibition of automatic breathing upon volitional control (Haouzi et al., 2006). Ventilation compensation by the Supplementary Motor Area (SMA) during arousal is suggested in patients with Congenital Central Hypoventilation Syndrome, who suffer from debilitating hypoventilation only during sleep (Tremoureux et al., 2014). Both previous examples illustrate complex interplay between volitional control and automatic commands of respiration, the former being able to temporarily bypass the later, an unique case for a life-sustaining physiological requirement in animals (Trevizan-Baú et al., 2024). Research in animals, further supported by one human neuroimaging investigation of volitional respiratory control (McKay et al., 2003) and a model of dyspnea (Betka et al., 2022), suggests a complex interplay between respiration-related cortical networks and the involvement of brainstem automatic respiration generation.

In the brainstem, a central respiration controller consists in a network of motoneurons located in the ventral reticular formation innervating the ribcage and upper airways (Viemari et al., 2013). Extensive animal studies has led to the identification of respiratory pattern generators, known together as the preBötzinger Complex (Smith et al., 1991), as central elements of this central controller. This complex orchestrates the three-phase - inspiration, post-inspiration and expiration - respiratory cycle (Bellingham, 1998). Because it is mainly devoted to controlling an automatic physiological reflex required for survival, respiration, it is not known how it contributes to voluntary control of breathing: actually, breathing may be an unique physiological reflex given it is usually automatic but can also be controlled, and in the case of speech, that requires to be synchronised with expiration, could be involuntary controlled but in conflict with the automatic control. Elucidating the respective role of cortical and brainstem substrates of respiration control in natural conversation is crucial to understanding how breathing can be associated either with survival or with communication in spoken social interaction.

### Current analysis

When we take turn during a conversation, the mechanisms discussed above harmonise in asymmetrical respiratory cycles (Rochet-Capellan & Fuchs, 2014). These cycles were measured as variations in lung amplitude and frequency over time using a respiration monitor belt, making it possible to monitor the various adaptations during a conversation. Significant differences can be observed between individuals depending on their age, gender and lifestyle, resulting in specific adjustments when breathing during speech. Studies have highlighted the influence of these factors on lung volume during spontaneous speech (Winkworth et al., 1995). Furthermore, within the same individual, intra-individual variability in lung volume during unscripted communication is also significant. Contextual factors, changing emotional states and particular cognitive demands contribute to this variability (Winkworth et al., 1995). These elements modulate the way a person adjusts lung volume during speech, creating a respiratory signature unique to each conversational context. This close coordination between the respiratory process and speech suggests that our respiratory system adapts dynamically to the demands of verbal communication, both inter-individually and intra-individually. It highlights the need to investigate this coordination taking into account sources of variability.

Here, we made use of a unique corpus of natural conversation comprising synchronised conversation, breathing and functional neuroimaging (fMRI) to contrast brain control of breathing according to whether it is associated with speech production (maximum occurring close to the participants’ speech onset) or physiological survival (necessity of pulmonary exchange between O_2_ and CO_2_). Conversation transcripts are used to categorise automatically breathing monitor belt maxima according to their temporal occurrence within a time window centred on the participant’s speech onset. Results indicate that cortical regions buried in the human central sulcus are significantly more active for speech-associated breathing maxima, as well as regions in the brainstem that we associate with human homologues of the respiratory rhythm generator known as PreBötzinger complex first identified in rodents (Smith et al., 1991). These results suggest that both cortical and subcortical control of breathing are involved in respiration control during turn-taking associated with natural conversation.

## Materials and Methods

### Corpus acquisition

Twenty-five native French-speaking participants (7 M, 18 F) with an average age of 28.5 years old (*s.d.* 12.4) were recorded in a fMRI protocol investigating natural conversations in Human-Human and Human-Robot Interactions. Experimental procedures were approved by the designated national ethical committee (“Comité de Protection des Personnes” CPP Sud-Marseille 1, approval number 2016-A01008-43) and respected all procedures in place. The conversational paradigm (Rauchbauer et al., 2019), its theoretical grounds (Chaminade, 2017) and the variables recorded (Rauchbauer et al., 2020), have already been described and will only be summarised. In particular, we will focus on the three variables utilised in the current analysis: brain activity (using fMRI Blood Oxygen-Level Dependent (BOLD) signals acquired continuously), respiration (focusing on inspiration-to-expiration transitions) and transcripts of the natural conversations (Hallart et al., 2021; Rauchbauer et al., 2020).

Participants were fully naive about the actual objectives of the current study, i.e. creating a multidimensional corpus of natural interactions including synchronised neurophysiological recordings. At the MRI centre were presented the cover story, that involved natural discussion about a purported neuromarketing experiment. During each functional scanning run, they underwent six trials alternating between conversation with a human and conversation with a robot. Each trial started with the presentation of one out of three images forming the neuromarketing cover story material, followed by one minute of live conversation with an interlocutor located outside of the scanner room. The bidirectional audio set-up enabled the live conversation between the scanner and the outside consisted of an active noise-cancelling MR compatible microphone (FORMI/III+ from Optoacoustics Ltd.) to reduce scanner noise and inserted earphones from Sensimetrics. Participants could see a live movie of the interlocutor projected on the scanner’s stimulation screen and had their eye movements recorded.

The interlocutor was either a confederate of the experimenter or the robotic head Furhat controlled by the confederate with a Wizard of Oz setup of ∼100 sentences prepared *a priori*. We recorded three trials each containing a one-minute block conversation per conversational agent (Human and Robot) in each scanning run (6 blocks total) and four runs per participant yielded an overall total of twenty-four minutes of conversation recorded. Effects of the conversational agent are irrelevant for the current analysis focusing on speech production mechanisms and won’t be discussed further.

The MRI data was collected with a 3T Siemens Prisma (Siemens Medical, Erlangen, Germany) using a 20-channel head coil. BOLD sensitive functional images were acquired using an EPI sequence in the four runs. Parameters were as follows: echo time (TE) 30 ms, repetition time (TR) 1205 ms, flip angle 65°, 54 axial slices co-planar to the anterior/posterior commissure plane, field of view 210 mm × 210 mm, matrix size 84 × 84, voxel size 2.5 × 2.5 × 2.5 mm^3^, with multiband acquisition factor 3. After functional scanning, structural images were acquired with a GR_IR sequence (TE/TR 0.00228/2.4 ms, 320 sagittal slices, voxel size 0.8 × 0.8 × 0.8 mm^3^, field of view 204.8 × 256 × 256 mm). Data acquisition of Blood Oxygen-Level Dependent (BOLD) signal 3-dimensional images were obtained via whole brain scans every 1.205 seconds. One run totalled 385 volumes for a duration of 463.925 s (7 minutes and 425 seconds). Raw data acquired during fMRI scanning have been uploaded on OpenNeuro^1^ including Text-format log-files containing trials onset information.

The physiological data (cardiac pulsation and respiration) in the study was recorded with the scanner’s dedicated apparatus. A photoplethysmography device was placed on the left-hand index fingertip to capture pulse oximetry, while a breathing belt was positioned at chest level. Both devices were connected wirelessly through Bluetooth, and the data was acquired continuously at a frequency of 200 Hz. Synchronisation is intrinsic to the data recording apparatus, and is based on the 2.5 ms time bins constitutive of MRI recordings: time is logged in bins for the BOLD signal and the breathing data, both recorded with the MRI scanner, and the conversational data processing also include a synchronisation and resampling step (see next).

### Data processing

fMRI preprocessing with SPM12 involved slice-timing corrections, volumes realignment and magnetic field inhomogeneities correction (Penny et al., 2007). Normalisation of individual functional and anatomical data to the standard brain space of the Montreal Neurological Institute (MNI) used segmented anatomical T1 and T2 images recorded at the just after BOLD functional images and realigned with the first image of the time-series, followed by the DARTEL procedure (Ashburner, 2007). BOLD signals were extracted from masks of the Grey matter, White matter and Cerebrospinal fluid individuals’ regions of interest derived from the segmentations to make detrending nuisance variable with the conn toolbox (Whitfield-Gabrieli & Nieto-Castanon, 2012), that incorporates other denoising procedures like linear detrending with a high-pass filter (threshold of 128 seconds) as well as realignment parameters for removal of artefacts due to participant movement calculated during the realignment step. Physiological recordings were processed using the PhysIO toolbox (Kasper et al., 2017) that employs model-based approaches for physiological noise correction of fMRI data. It uses physiological measurements to derive heart rate variability (referred to as the cardiac response function) and respiratory volume per time (referred to as the respiratory response function). Both movement-related and physiology-related possible sources of artefacts were exported as single-participant and single-run matrices containing aggregating cardiac, respiration and movement signals possibly associated with recording artefacts as well as linear and region-of-interest detrending nuisance variables. Finally, we applied a 5 mm^3^ full-width half-maximum three-dimensional Gaussian kernel spatial smoothing.

The respiration time-series were processed in order to ensure signal integrity by replacing plateau signals corresponding to saturation of the recording belt signal. The respiration data was first smoothed using a moving average with a window size of 50. Regions of saturated signal were identified by contiguous segments where the derivative of the breathing time-series was null. Consecutive numbers in these segments are replaced by ‘NaN’ so that saturation is considered to represent missing data. The missing data were replaced by non-linear interpolations (*spline* method in MATLAB® R2022a’s function *interp1*) taking into account the full duration of each individual run. Time series were finally resampled in milliseconds synchronised with fMRI acquisition, taking advantage of the fact that the offset of the respiration time-series is synchronised with the end of the fMRI recordings.

Conversational data from participants’ audio recordings underwent preprocessing to eliminate noise during scanner acquisition, employing a noise reduction filter provided by Sox (http://sox.sourceforge.net). For each participant, an individual float value, ranging from 0.01 to 0.50, was assigned. The denoised data was subsequently segmented into Inter-Pausal Units (IPUs) with each participant having an individual coefficient, a float value between 0.20 and 0.95. This coefficient facilitated the automatic determination of volume thresholds for periods classified as “silence” and “IPU” within each analysis window. It was applied to the mean of the root mean square (RMS) distribution of the audio file. IPUs were defined as speech blocks occurring between silences, with minimum duration of 200 ms (Blache et al., 2009). Visual validation of successful denoising and IPU segmentation was conducted using Praat. Files were furthermore uploaded into SPPAS, version 1.9.9 (Bigi, 2015) for transcription. Automatic Text normalisation was performed using the SPPAS software tool.

### Definition of experimental events

Automatic processing algorithm were developed to 1) identify local respiration maxima as they correspond to transitions between inspiration and expiration associated with neuronal events (Bellingham, 1998) that we expect to differ as a function of 2) whether or not single maximum are temporally associated with speech (operationalized as IPU) onset.

First, we used the log files related to fMRI recording to identify single block onsets, then added this value to the onsets of participant’s produced IPU in this block (i.e. the moment the participant takes its turn of speech in the conversation) to obtain for each run across blocks the onset of each speech-turn from the scanned participant during the conversations. The respiration maxima temporally closest (in absolute value) from every IPU onset was categorised as “*Respiration maxima associated with speech production*” (Resp+), while all others were attributed to “*Respiration maxima not associated with speech production*” (Resp-). Attribution of single Resp+ events to every IPU event allowed us to calculate the relative time difference between these two events Δt.

### Analyses

Analyses were performed across the 4 runs of the 25 participants included in the corpus. For the behavioural analysis, we used MATLAB® R2020a’s function *rmoutliers* to identify and remove outliers from Δt values, i.e. values more than three median absolute deviations from the median. The missing data is below 5%, allowing the use of all data in the analysis. Kolmogorov-Smirnoff tests of normality were run on the resulting cleaned Δt values on the final time values distribution and reached the significance level (*p* < 0.05) for most participants and runs, indicating that the data is not normally distributed. We therefore performed skewness tests to measure the asymmetry of the distribution.

For each participant and run, BOLD data was analysed with SPM12 with three conditions, Image presentation, that will not be discussed furthermore, and the two conditions of interest, Resp+ and Resp-modelled as events (*i.e.* zero duration). All nuisance variables calculated with the PhysIO toolbox were introduced in these models, allowing to remove artefacts associated with physiological (breathing and heartbeat) noise, movement artefacts and detrending based on linear as well as regions-of-interest signal (Kasper et al., 2017).

Given that Resp+ events are temporally synchronised with IPUs onset, therefore hindering the distinctions between respiration and articulatory correlates of speech, we used an exclusive masking procedure to remove brain activity associated with speech production in our corpus. To do so, we performed another analysis with conditions “*IPUs from participant*” (IPU+) and “*IPUs from interlocutor*” (IPU-), describing each produced and perceived speech turn by its onset and its duration. The activation map corresponding to the contrast IPU+ *versus* IPU-, assessed at the second level (with the same parameters as in the main analysis presented in the next paragraph; *p_FWE_* < 0.001) was saved as a binary image, and expanded by two voxels in all directions (using MRIcron), to avoid border effects, to create a mask including brain areas involved in articulatory aspect of speech.

For the second-level analysis of interest, individual beta estimates from the two conditions of interest Resp+ and Resp-were used in a full ANOVA model using SPM12, with participants and runs as repeated measures (unequal variance for the former, equal for the later), using the mask described previously to exclude from analysis brain regions associated with speech production. Results for the contrast Resp+ versus Resp-are provided with a threshold of *p_FWE_* < 0.05 at the peak level and *p_FDR_* < 0.05 at the cluster level (corresponding to an extent k > 5 cm^3^).

### Central sulcus localization

We conducted a single subject and single run analysis to identify more precisely the location of the central sulcus cluster in each hemisphere, in particular to distinguish between precentral and postcentral localizations. Significant activity (*p_unc_* < 0.001) was found within a 1 cm radius around the group analysis peak in 145 out of 200 possibilities (25 participants x runs x 2 hemispheres) *i.e.* almost 3/4. We developed a procedure to minimise subjective biases to localise the respective position of the peak of individual activity in respect to the position of the central sulcus. Human intervention was minimised to two aspects of the procedure, while automatic MATLAB® and python scripts were used to draw the central sulcus and overlay the central sulcus drawing on the activity map for each participant, run and hemisphere in which activity was found.

We first saved an axial section of the normalised structural image made at the height of the peak of the identified cluster, usually around z = 60 mm in MNI coordinates, without any activity overlaid. Three independent observers traced the central sulcus on these images. The absence of any discrepancy between the observers validated the identification of the central sulcus on each structural section. The images were then cropped manually with Photoshop to contain only the central sulcus region identified. The tracing of the central sulcus was semi-automated using *ad hoc* python scripts that draw the sulcus based on the grey value intensity of the anatomical image. To bootstrap sulcus drawing, a starting point was selected manually. As the sulcus never reaches the midline at this level of the brain, we identified the pixel that could still be identified as grey matter found at the fundus of the central sulcus closest to the midline. A python script draws a line corresponding to the central sulcus iteratively by looking for the adjacent pixel with the lowest intensity centrifugally.

Finally, a MATLAB® script overlaid the line drawing corresponding to the central sulcus to the functional image in which the peak location of the activated cluster is indicated by a cross yielding a series of images allowing us to categorise the respective location of the individual activation peaks and the central sulcus., with four possibilities: 1) the cross falls on the line, 2) at the fundus when it is close (less the one pixel) from the starting point chosen arbitrarily, and for all others, 3) anterior or 4) posterior to the central sulcus.

## Results

### Behavioural analysis

Analysis of skewness in Δt revealed a significant left-skewed distribution, with a maximum around −200 ms (-0.2003 ms) illustrated on Figure 1. This indicates that a maximum breathing intake occurred on average 200 ms before participants speech onset. The finding of maximum breath following speech onset (in red on Figure 1) could be surprising, but it could be due to timing errors associated with both the breathing maximum estimate, as interpolation of saturated signal could fail to reproduce some aspects of the breathing signal such as the asymmetry between the dynamics of inspiration and expiration (Winkworth et al., 1995). Additionally, timing of IPU onset also suffers from possible timing errors depending on the type of phonemes produced at the beginning of the speech production. Importantly, while these are intrinsic limitations of the current investigation, they are unlikely to interfere with fMRI results given its poor temporal limitation. the statistical power of the fMRI analysis comes from the repetition of events of interest, which combine all respiration cycles recorded during the one acquisition run, *i.e.* several hundreds of breathing maxima for an approximate 7-min & 45-sec runs, with variance in the timing between events related to other behavioural factors such as the presence or absence of speech.

**Figure 1.**
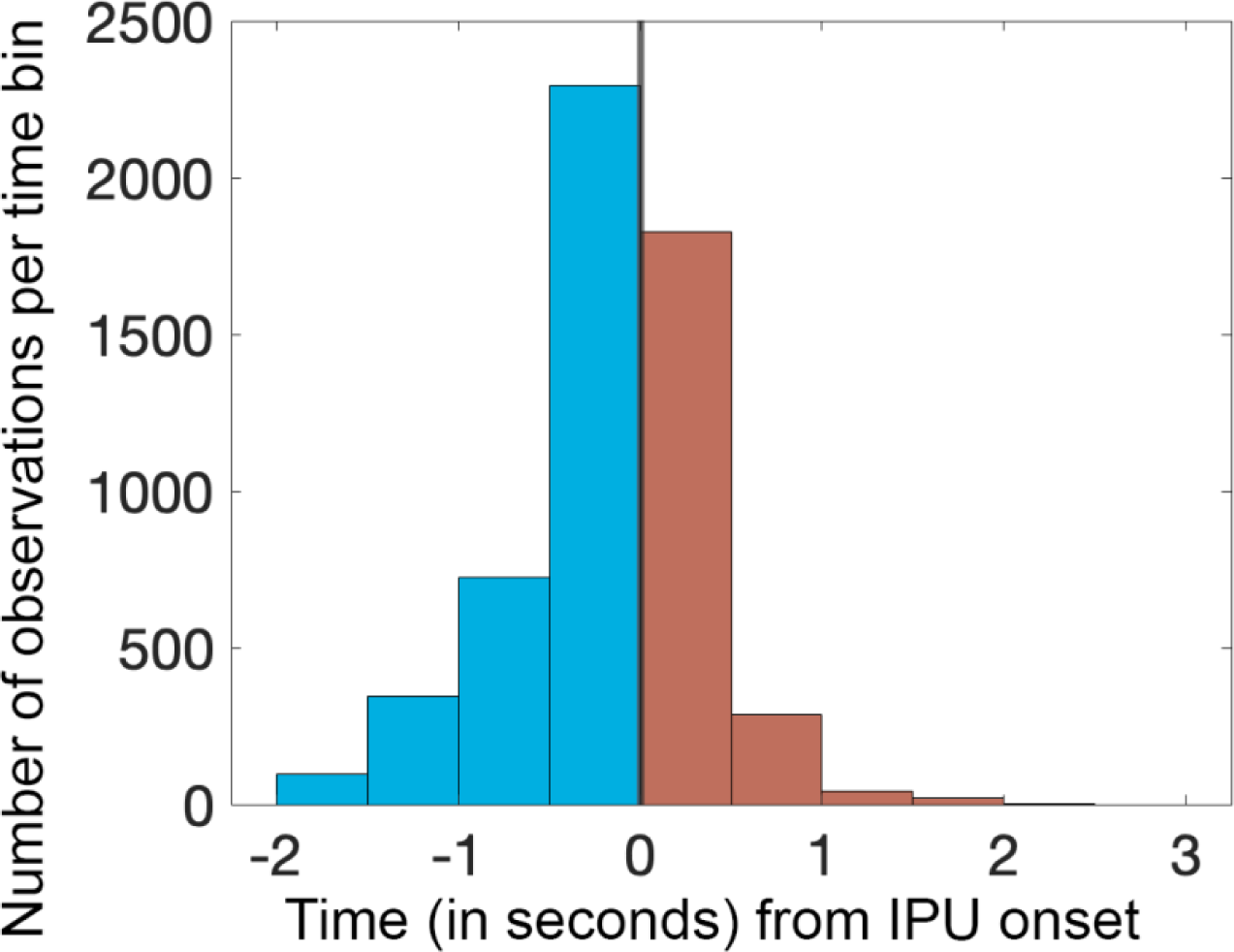
Time difference Δt between IPU onset and max breathings indicates a left skewness peaking at 200 ms, indicating that maximum of breath occurred on average 200 ms preceding participants speech onset.

### Second level analysis of respiration maxima related to IPUs

Second level contrast Resp+ *versus* Resp-(*p_FWE_* < 0.05, extend k > 5 cm^3^) exclusively masked with the contrast IPU+ *versus* IPU- (*p_FWE_*< 0.001) reveal bilateral activations in the central sulcus, in the brainstem and in the dorsal and ventral cerebellum (see Table 1 and Figure 2). Note that the reverse contrast revealed no activations at the thresholds used. Attribution of cerebellum clusters to the primary fissure and hemispheric lobule VIII for the dorsal and ventral clusters respectively relied on a cerebellum atlas allowing the use of MNI coordinate system with visual inspection of the cluster with regards to cerebellar folia boundaries (Schmahmann, 2000).

**Figure 2:**
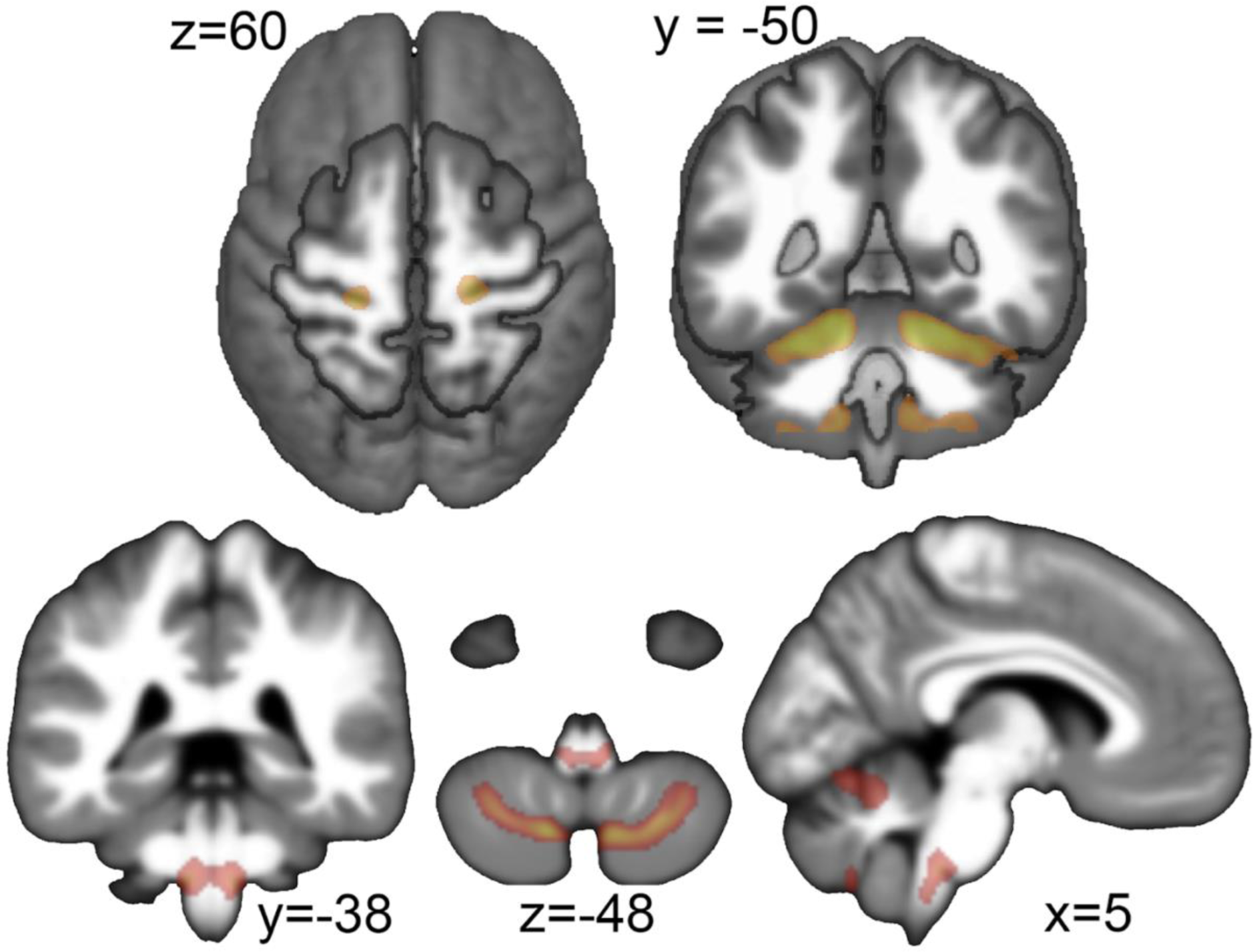
Brain activations for maximum breath intake associated with a speech onset (Resp+ *versus* Resp- [*p_FWE_* < 0.001, extend > 5 cm^3^] exclusively masked by IPU+ *versus* IPU-). Top left: the bilateral central sulcus fundus clusters. Top right: Dorsal and ventral (also see bottom, centre) cerebellar bilateral clusters. Bottom: coronal, axial and sagittal views of the brainstem cluster.

**Table 1.** Significant activations associated with the contrast Resp+ *versus* Resp- (*p_FWE_* < 0.001, extend > 5 cm^3^). MNI coordinates (in mm) are used to locate the clusters of significant increased activity.

| *Location* |  | *Extend*<br>*(voxels)* | *peak*<br>*T-values* | *MNI coordinates (mm)* |  |  |
| --- | --- | --- | --- | --- | --- | --- |
| *Anatomy* | *Side* |  |  | *x* | *y* | *z* |
| *Central Sulcus*<br>*(fundus, dorsal part)* | *R* | *819* | *16.13* | *20* | *-27* | *60* |
|  | *L* | *681* | *15.28* | *-21* | *-30* | *63* |
| *Cerebellum*<br>*(centred on Primary Fissure)* | *R* | *5676* | *21.36* | *14* | *-60* | *-20* |
|  | *L* | *\** | *20.96* | *-15* | *-60* | *-18* |
| *Cerebellum*<br>*(hemispheric lobule VIII)* | *L* | *763* | *14.38* | *-12* | *-66* | *-50* |
|  | *R* | *1141* | *14.16* | *10* | *-68* | *-48* |
| *Brainstem*<br>*(ventral reticular formation)* | *L* | *706* | *10.82* | *-8* | *-38* | *-45* |
|  | *R* | *\** | *10.47* | *8* | *-38* | *-45* |

The precise attribution of the respiration clusters to pre- or post-central gyrus, triggered by the observation of Figure 2 (top left), was analysed and results are illustrated in Figure 3 (single participant and runs images are provided in Supplementary figure 2). Although these results suggest that clusters maxima are mainly found in the postcentral gyrus, a comparable number of maxima were attributed to the anterior or fundus region of the central sulcus. But it has been reported that anatomical (the fundus of the sulcus) and functional (transition for primary sensory to primary motor cortices) don’t match and the boundary can extend towards the anterior precentral gyrus (see *e.g.* Fig. 10 in White et al., 1997). By extrapolation, considering that all postcentral gyrus clusters should be attributed to the primary sensory cortex and even a fraction of the clusters associated with the other cortical regions could also belong to this functional area, we propose that the bilateral central sulcus activity reflects a neural correlate of sensory processing related to Inspiration-to-Expiration transitions.

**Figure 3:**
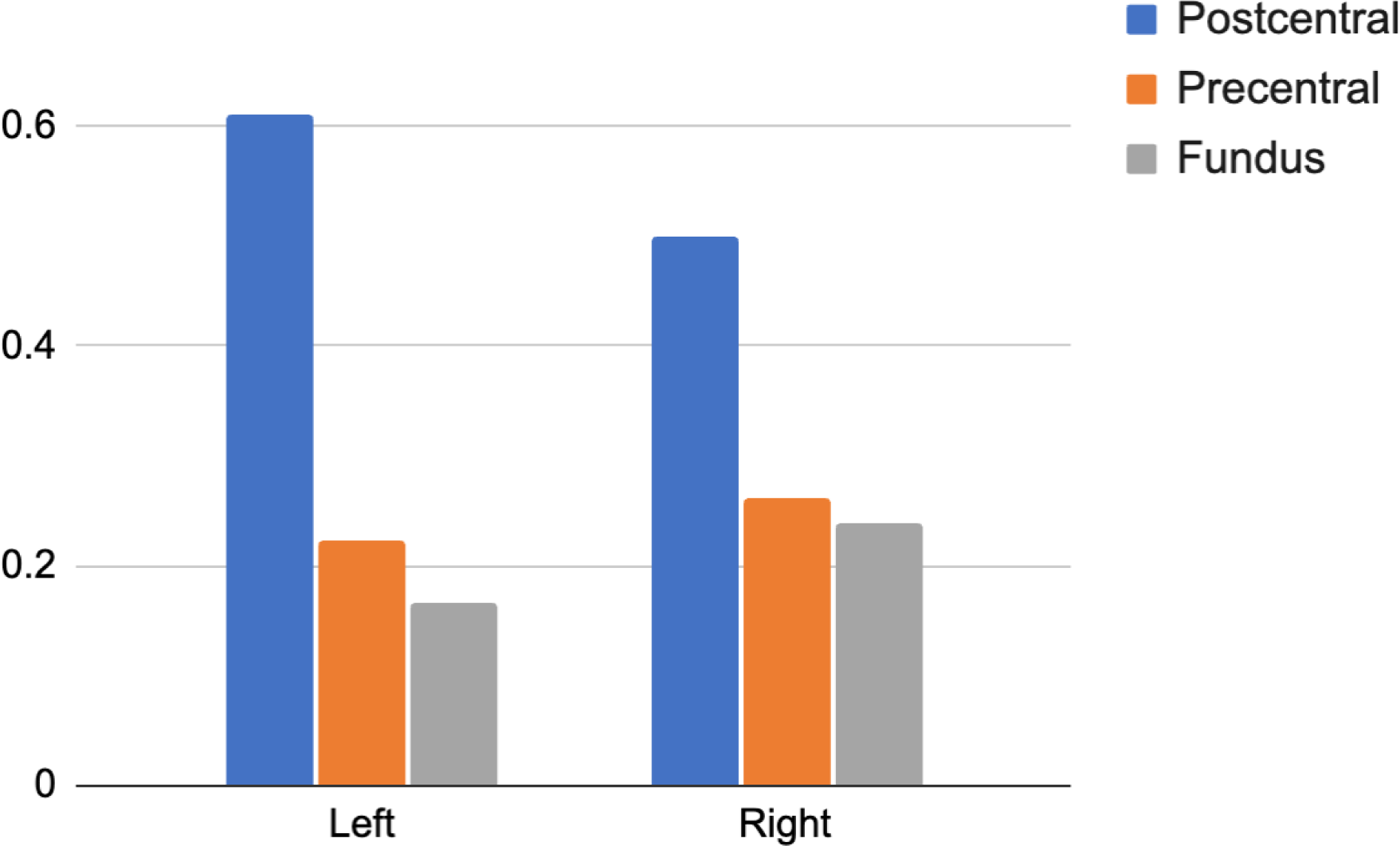
Individual participants and sessions contrast Resp+ *versus* Resp- (p_unc_ < 0.001) within a 1-cm radius sphere centred on the group central cluster peak of activity were semi-automatically attributed to the fundus of the sulcus and the pre- and the post-central gyri. The figure presents the number of observations for the three possible cortical assignments.

Finally, localisation of functional areas in the brainstem is particularly challenging given the small size of brainstem nuclei compared to the fMRI voxel size, uncertainties about interindividual variability and the absence of a reference atlas. A post-mortem brainstem atlas indicated that the brainstem cluster is located between the cerebellocortical and thalamocortical bundles identified with tractography (Adil et al., 2021). Circumscribed by white fibre tracts and spanning medullary and pontine regions, this bilateral cluster, elongated along the main axis of the brainstem identified with a contrast investigating maximum respiration events, likely belongs to the human ventral reticular formation where is located the homologue of the pre-Bötzinger complex identified in rodents (Smith et al., 1991).

## Discussion

The aim of the present study is to investigate the neurophysiological underpinnings of speech-related respiration events in conversational turn-taking. We analyse an existing corpus of natural conversations between a participant and its interlocutor (Human or Robot) focusing on synchronised (1) behavioural (conversation turn-taking), (2) respiratory (maxima of inspiration and expiration) and (3) neurophysiological (fMRI) recordings. Each participant’s speech production during the conversation was recorded, denoised and segmented into silence and speech turns. Participants’ speech turns (or InterPausal Units, IPUs) were defined as speech blocks occurring between silences, each lasting at least 200 ms (Blache et al., 2009).

### Respiration maximum and speech onset

We first investigated the temporal relation between respiration maxima and speech production. The closest respiration maximum, in absolute value, was associated with each IPU, hence referred to as “Respiration maxima associated with IPUs” or “Resp+”. We further investigated the distribution of temporal associations between expiration and vocal production by focusing on the relative time differences between the two categories of events. As illustrated in Figure 1 with the cumulative histogram across all participants, runs, trials and turns indicated that maximum respiratory intake occurred on average 200 ms prior to speech onset. The variability could be due to the accumulation of temporal noise, both from the fMRI recordings precluding millisecond precision in the identification of participant’s speech onset and from the saturation of the recording belt signal that occasionally required extrapolation. Nevertheless, Supplementary figure 1 shows that the blue bar left to the 0 ms (interval [-500; 0] ms) has the highest number of observations for most participants and runs.

This result aligns with studies indicating a temporal gap between the cessation of air intake (respiration maximum) preceding the onset of speech by 200 ms (MacIntyre & Scott, 2022). Other studies have considered various factors. For example, one study found that inhalation depth correlates with the speech envelope’s total power and relates to upcoming speech, suggesting internal models for enhanced communication (Abbasi et al., 2023). Another study highlighted that respiration and pauses are crucial for turn-taking, showing that successful turns often occur right after a new inhalation, indicating that speakers coordinate their breathing with turn-taking (Rochet-Capellan & Fuchs, 2014).

Altogether, the temporal relationship between speech and respiration appears to be preserved despite the lack of naturalness experienced when speaking while lying supine in a scanner during a noisy fMRI acquisition. For the fMRI analysis, we defined “Resp-” events as “Respiration maxima not associated with IPUs”. Together, Resp+ and Resp-identify all respiration maxima, and are used as events for the fMRI analysis. Note that the physiological data (cardiac and respiratory cycles) were processed using the PhysIO toolbox (Kasper et al., 2017) and used to derive nuisance variables describing noise induced by heart pulsation, respiratory volume as well as individual participants’ movements derived from functional magnetic resonance images (fMRI) realignment. Physiological artefacts being removed with this denoising procedure, the results obtained reflect central nervous system respiratory control associated with conversational speech onset and not artefacts derived from peripheral physiology (Raitamaa et al., 2021).

The physiological aspects of speech breathing have shown muscular components, represented in the motor cortex, that contribute to the regulation of breathing required for voluntary control and sound production (McKay et al., 2003; Winkworth et al., 1995), underlining its clear differentiation from metabolic breathing. The unique data used here, a natural conversation, allows us to further the understanding of breathing control in a social interaction. To identify brain correlates of speech-related breathing maximum, we compared events described as Resp+ to Resp-using a classical preprocessing of data and two-stage analysis in SPM12 (Penny et al., 2007). As this contrast yielded results clearly associated with speech production (in premotor cortices in particular), a masking procedure was used. A mask describing the main effect of speech production was created from an independent analysis of the same data defining each IPU as a block to identify. Using this mask to exclude speech associated brain regions at the population level, allowed us to identify areas significantly more activated in the contrast Resp+ *versus* Resp- (p_FWE_ < 0.001, extend k > 5 cm^3^) and not associated with speech production. This analysis revealed bilateral activations in the dorsal and ventral cerebellum, in the brainstem and in the central sulcus, all these clusters being further described in Table 1 and illustrated in Figure 2.

### Speech-associated respiration maxima and central sulcus response

Several regions around the central sulcus are involved in modulating respiratory activities. Studies have shown that specific regions are activated during the initiation and coordination of voluntary respiratory movements. These regions send signals to brainstem structures involved in regulating respiratory rhythm such as the inferior olive bilaterally (Sears et al., 1995). The central sulcus is also involved in speech production. Specifically, the primary motor cortex is responsible for the control muscles necessary for producing speech sounds. It sends signals that are transmitted through regions of the brainstem (though the exact pathways remain elusive, see Belyk et al., 2021) to control precise movements of the laryngeal muscles, vocal folds, and articulations involved in speech. Motor cortices are therefore essential in coordinating and regulating speech-producing movements.

The exact location of the cluster at the fundus of the central sulcus (see Figure 2) made it difficult to attribute the response across participants to the pre- or post-central gyrii, especially given that the boundary between cytoarchitectonic Brodmann Areas 3 (primary somatosensory cortex in the postcentral gyrus) and 4 (primary motor cortex in the precentral gyrus) is at the fundus of the central sulcus (White et al., 1997). Hypothesising that the data smoothing associated with DARTEL normalisation and group-level analysis could explain the impossibility of a clear attribution, we analysed the localization of the cluster in single participants and single runs with unsmoothed anatomical and functional data in a semi-automatic fashion. Results showing the relative position individual participants’ cluster of activity in the central sulcus drawn (shown in Supplementary figure 2 and summarised in Figure 3) reveal that a majority of activated clusters for the contrast Resp+ *versus* Resp-are found in the post-central sulcus. As this would locate them in primary sensory cortices, the central sulcus cluster would be related to sensory processing, and not motor control. While speculative, these results suggest that sensory information from the chest upper body area, for example the diaphragmatic aspects of respiratory functions, could be processed in this primary sensory region.

### Speech-associated respiration maxima response in the brainstem

This sensory component found in the central sulcus might be associated with “perceiving” the maximum intake of oxygen in the lungs required for producing subsequent speech. This could in turn also explain increased response for Resp+ events in brainstem regions associated with control of breathing. The cluster was attributed to the ventral respiration controller preBötzinger complex, which contains both inspiration and expiration neurons. This complex has multiple subcortical, but very limited cortical, projections in rodents (Yang & Feldman, 2018).

The main contribution of preBötzinger complex is to generate rhythmic respiratory patterns (Smith et al., 1991). Studies suggest that connections indeed exist between the motor cortex and brainstem regions involved in respiratory regulation that include the preBötzinger complex as well as other interconnected pontine and medullary nuclei such as the nucleus ambiguus, the inferior olive or the periaqueductal grey matter also involved in vocal production (Zhang & Ghazanfar, 2020). These connections enable bidirectional communication between the cortex and the brainstem to coordinate respiratory movements necessary for speech production (Belyk et al., 2021). The interactions may occur through descending projections from the motor cortex to the brainstem, regulating preBötzinger complex activity and influencing respiratory control. Reciprocally, signals from several brainstem nuclei project back to the cortex and can carry sensory information about the current phase of the respiratory cycle given the level of inspiration.

### Speech-associated respiration maxima and cerebellar response

This temporal dynamic is reflected in the third result of the fMRI analysis regarding the attribution of cerebellum clusters in the bilateral primary fissure (V, VI) and right hemispheric lobule VIIIa and VIIIb more activated in the condition Resp+.

The cerebellum, a brain structure traditionally associated with motor coordination and balance, has increasingly drawn attention for its significant role in speech production and perception (Ackermann et al., 2007). They suggest that the cerebellum plays a crucial role in regulating speech rate, with evidence indicating its involvement in pushing speech rates beyond 3 Hz, emphasising its importance in the temporal organisation of verbal utterances. Neuropsychological studies confirm its involvement in various speech-related conditions, such as agrammatism and transcortical motor aphasia in patients (Ackermann et al., 2007). Transcranial stimulation of the right cerebellum improved naming abilities in individuals with aphasia (Sebastian et al., 2020). Moreover, ataxia, a condition characterised by cerebellar impairment, has been associated with significant deficits in spontaneous speech reducing intelligibility and naturalness (Hilger et al., 2022). In addition, damage to cerebellar fibre tracts in the lobules VI of the cerebellum may further compromise speech motor control, contributing to ataxic dysarthria, a speech disorder characterised by difficulty in articulation and pronunciation (Lehman et al., 2020). Other studies highlight the functional convergence of motor and social processes in lobule IV/V, suggesting a broader role in cognitive and social behaviours (Chao et al., 2021). Lobule VIII is implicated in language processing, with studies reporting correlations between greater grey matter volume in lobules VII and VIII and better performance on measures of language (Stoodley & Schmahmann, 2018).

Importantly, one function repeatedly associated with the cerebellum is the timing associated with motor control (Tanaka et al., 2021). Given the importance of the relative timing between the events under investigation, the maximum respiration intake, and the relative timing of a specific action, speech onset, it is proposed that the cerebellar responses that we report associated with maximum respiratory events are linked to the preparatory components of vocal production.

### A cortico-brainstem-cerebellar network for speech turn-taking in natural conversation

We therefore have three actors involved in the coordination between respiratory maximum and speech onset during natural conversation: the posterior part of the central sulcus dorsally, a region of the brainstem likely to be associated with brainstem respiratory pattern generator, and regions of the cerebellum that could be involved in temporal coordination of motor control. Increased response in the brainstem cluster, supposedly involved in the generation of breathing rhythms, likely reflects the active inhibition of the respiratory pattern generators that is required to enslave expiration to vocal production. Had this inhibition been passive, we would have expected an increased response in the brainstem in the opposite contrast, focusing on maximum respiratory intake not associated with speech production, which yielded no significant response in the current analysis. The source of this inhibition could be the cerebellum clusters found in the same contrast, as it has been reported a direct inhibition of respiration by electric stimulation of Purkinje cells (reviewed in Xu & Frazier, 2002). Such interpretation is in agreement with the role of the cerebellum in the temporal aspects of motor control. Finally, the central sulcus cluster, attributed to the primary sensory cortex, could detect by afferents from upper airways the moment when the lungs are full and therefore ready to initiate speech. This important piece of information could also be integrated by brainstem and/or cerebellar components. Altogether, the relatively complex, but fully automatic, synchronisation of breathing with speech production involves the combination of several information reflected in the activation of regions belonging to different regions of the central nervous system.

## Conclusion

Our study confirmed a temporal dynamic between respiration and conversational dynamics, and identified several key actors involved in this dynamic at different levels of the central nervous system. These actors are the dorsal central sulcus, possibly corresponding to the primary sensory cortex encoding information coming from the chest level that can be associated with the filling of the lungs, the cerebellum, likely involved in computing the temporal dynamic between lung filling and speech onset, and the brainstem, where respiratory rhythm generators of the preBötzinger complex located in the ventral reticular formation must be inhibited to enslave respiration to speech production. While speculative, these interpretations are based on a large but parcellar literature that investigated the different actors in isolation or by pairs, especially in animal studies, complemented by neuropsychological results of speech impairments in humans reviewed in introduction. But to our knowledge, it is the first time that a corpus of natural conversation by humans including synchronised functional neuroimaging is used to identify the actors in relation to breathing and speech turn-taking. While further research will be necessary to investigate the dynamics of this central nervous system network, these results underscore the importance of integrating peripheral physiological signals to investigate complex behaviours in ecological settings.

## Acknowledgements

CDP is supported by the Institute for Language, Communication and the brain (ILCB ANR-16-CONV-0002). The research is supported by grants ANR-16-CONV-0002 (ILCB), ANR-11-LABX-0036 (BLRI) and AAP-ID-17-46-170301-11.1 by the Excellence Initiative of AixMarseille University (A*MIDEX), a French “Investissement d’Avenir” programme and the Institute for Language, Communication and the Brain.

## Footnotes

1 https://openneuro.org/datasets/ds001740

## Notes

### Competing Interest Statement

The authors have declared no competing interest.

