## Supplementary figures and images for "Central Nervous System Control of breathing in Natural Conversational Turn-Taking"

### Supplemental figures 1 and 2

#### Supplementary Materials

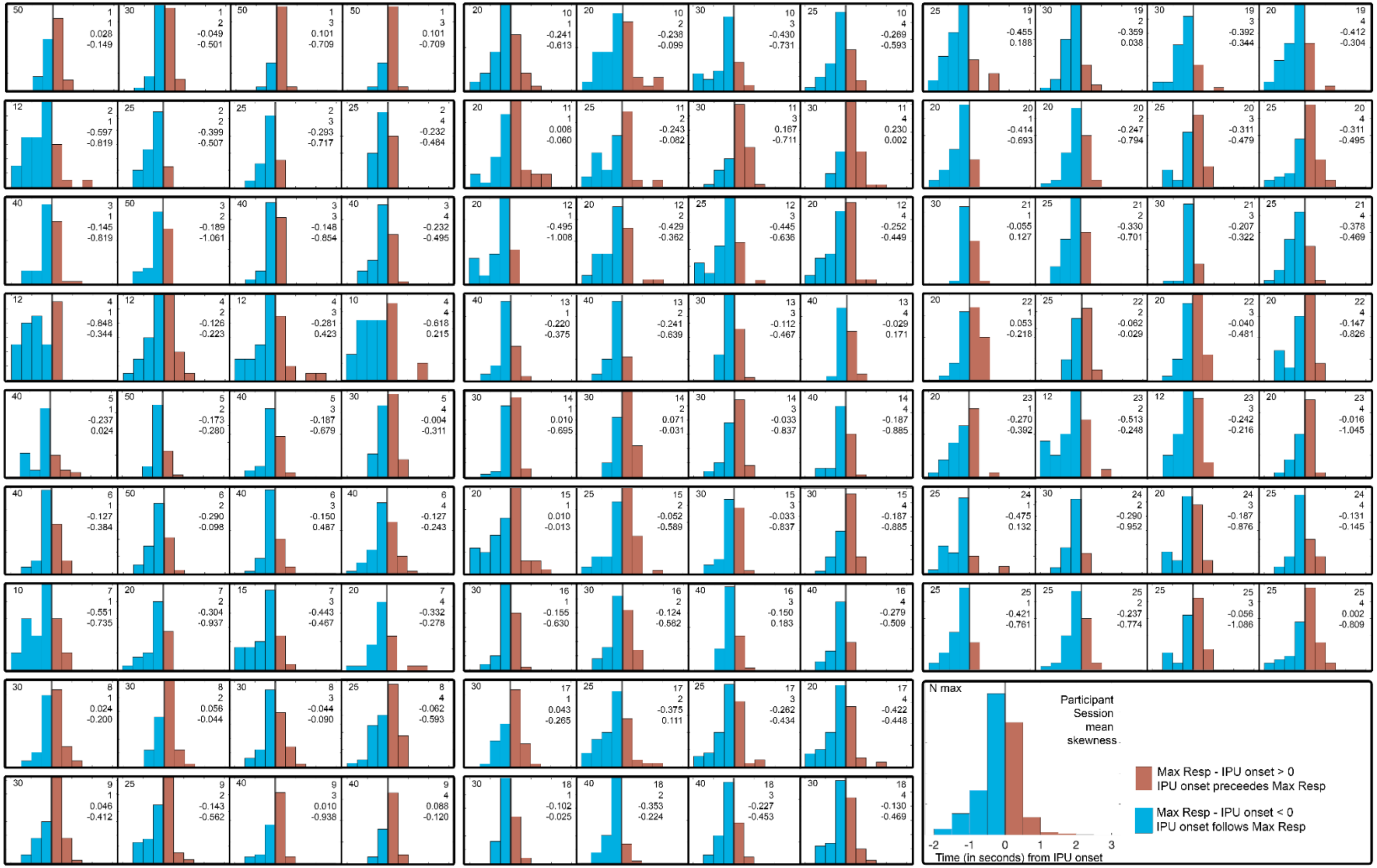


**Supplementary figure 1**

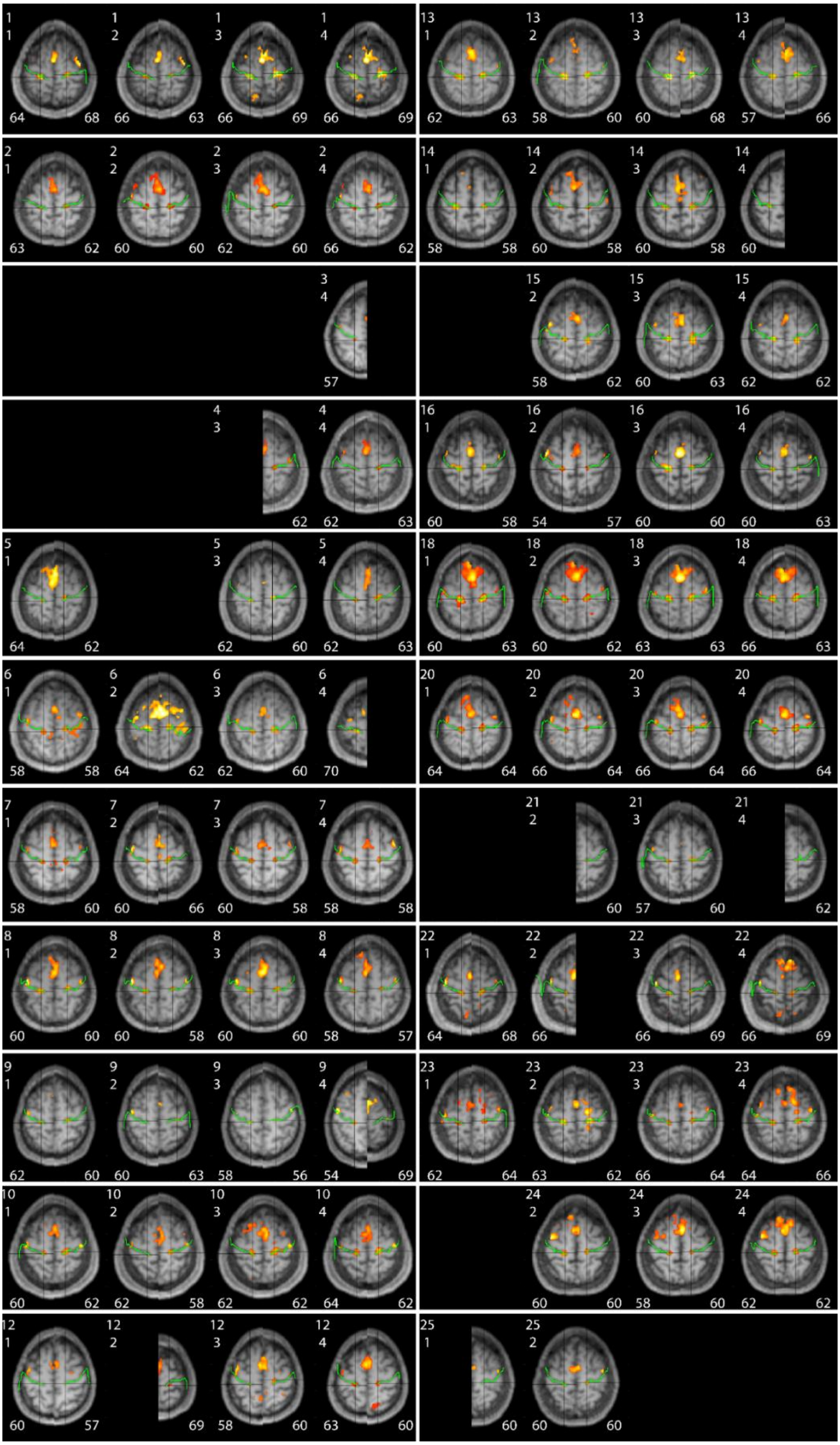

**Supplementary figure 2**
